# Computational Identification of Ligand-Receptor Pairs that Drive Human Astrocyte Development

**DOI:** 10.1101/2022.05.31.491513

**Authors:** AJ Voss, SN Lanjewar, MM Sampson, A King, E Hill, A Sing, C Sojka, SA Sloan

## Abstract

Extrinsic signaling between diverse cell types is crucial to nervous system development. Ligand binding is a key driver of developmental processes, but it remains a significant challenge to disentangle how collections of these signals act cooperatively to affect changes in recipient cells. In the developing human brain, cortical progenitors transition from neurogenesis to gliogenesis in a stereotyped progression that is influenced by extrinsic ligands. Therefore, we sought to use the wealth of published genomic data in the developing human brain to identify and then test novel ligand combinations that act synergistically to drive gliogenesis. Using computational tools, we identified ligand-receptor pairs that are expressed at appropriate developmental stages, in relevant cell types, and whose activation is predicted to cooperatively stimulate complimentary astrocyte gene signatures. We then tested a group of five neuronally-secreted ligands and validated their synergistic contributions to astrocyte development within both human cortical organoids and primary fetal tissue. We confirm cooperative capabilities of these ligands far greater than their individual capacities and discovered that their combinatorial effects converge on AKT/mTOR signaling to drive transcriptomic and morphological features of astrocyte development. This platform provides a powerful agnostic framework to identify and test how extrinsic signals work in concert to drive developmental processes.

**HIGHLIGHTS:**

- Computational prediction of active ligand-receptor pairs in the developing brain
- Synergistic contributions of predicted ligands drive astrocyte development
- Ligands induce transcriptomic and morphological features of mature astrocytes
- Cooperative ligand activity converges on AKT/mTOR signaling

## INTRODUCTION

Nervous system development involves interactions across diverse cell types in both time and space^1^. Each stage of this process must be carefully choreographed, from the temporal dynamics of cell fate commitment^2^ to the spatial migration and connectivity of developing circuits^3^. Throughout development, crosstalk among and between different cell types is essential and provides a mechanism that can influence cell state, identity, migration, pathfinding, and connectivity.

The dynamic interactions and crosstalk between neuronal and glial cells are particularly crucial for proper brain development^4–7^. Neurons and astrocytes share a common neuroepithelial origin and are born throughout embryogenesis in a temporally-defined manner^8–10^. The first divisions of neural stem cells called radial glia (RG) are exclusively neurogenic, either giving rise to neural-restricted intermediate progenitors or directly to young neurons^11^. Once the bulk of neurogenesis is complete, radial glia transition to a primarily gliogenic fate, and this change is referred to as the gliogenic switch^10^.

Both extrinsic and intrinsic signals can influence the gliogenic switch and the fate commitment of RG during development^8,12–19^. For example, several key transcription factors including NFIA^17,18,20^, SOX9^18,21,22^, RUNX2^23^, and RORB^24^ have been implicated as drivers of astrogenesis, and these intrinsic signaling pathways are either activated by extrinsic cutes^25^ or act synergistically with soluble ligands^26^ to affect astrocyte development.

Additional evidence for the role of extrinsic signals in the onset of astrogenesis arose from the observation that mouse embryonic radial glia produce neurons when cultured on embryonic cortical slices, but shift towards a glial fate when cultured on postnatal cortical slices^27^. Several secreted cues and membrane-bound signals have subsequently been implicated as gliogenic effectors, most of which belonging to the IL-6, BMP, and Notch signaling families^8,28–30^. The IL-6 subfamily of molecules (CNTF, LIF, CT-1, NP and CLC) all act via the signal-transducing coreceptors LIFRβ and gp130, and mice lacking either *LIFR*β *or gp130* have deficits in astrogenesis^12,28,29^. Other cues like BMP4^31–35^ and TGFβ^36^ family members influence astrocyte development through STAT3^16,37,38^ and SMAD^39^ activation, which can bind directly to astrocyte targets like the GFAP promoter^37^. Importantly, many of the extrinsic factors that have been identified thus far to influence astrocyte development are thought to be secreted from newly-born cortical neurons^10,14,27,40^. This creates an inherent timing mechanism whereby the extrinsic cues that are required for astrocyte formation may be supplied by the neurons whose development immediately precedes gliogenesis.

Importantly, while great strides have been made to identify individual ligands that exhibit gliogenic capacity, neural progenitors *in vivo* are simultaneously exposed to a varied assortment of extrinsic cues. Whether these signals work synergistically to influence cell fate has been difficult to determine given the enormous potential combinations among thousands of known secreted ligands. This is further complicated by the fact that the potency of extrinsic cues may depend on: (1) the intrinsic receptivity of RG to these ligands, (2) local ligand concentrations, and (3) their ability to coordinate activation of a broad signature of astrocyte genes. Given that individual ligands have been shown to have important contributions to astrocyte development^32,33,38^, we hypothesized that synergistic effects of specific groups of extrinsic signals may exhibit even more profound impacts on gliogenesis.

Here, we leverage existing bulk and single cell data to computationally predict novel combinations of ligand-receptor pairs that influence human astrocyte development. While all CNS cell types are likely to have distinct contributions to astrocyte development, we focused this study on neuronally derived signals, since these are among the most abundant cells in the brain at the time of astrocyte formation^2^. To identify cohorts of candidate ligand-receptor pairs that likely influence astrocyte development, we applied input data from a series of existing mouse and human transcriptomic datasets into an *in silico* framework called NicheNet^41^. This approach has the added benefit that it incorporates prior models of intracellular signaling to prioritize candidate ligand-receptor pairs by their potential to modulate complementary astrocyte gene signatures. We then apply a suite of transcriptional, morphological, and protein phosphorylation assays to demonstrate that combinatorial exposure of 5 specific ligands (TGFβ2, BMP4, DKK1, TSLP, and NLGN1) promotes astrocyte development in both an *in vitro* human cortical organoid model as well as primary human fetal astrocytes. In all assays, we observe that synergistic application of our ligand cocktail exhibits effects on astrocyte development that eclipse individual effects of each ligand. Additionally, we identify specific temporal windows of RG receptivity to gliogenic ligands and use protein-level readouts to assay candidate signaling pathways that drive astrocytic responses in the presence of the gliogenic cocktail.

Altogether, this study provides a powerful framework to apply existing datasets towards the discovery of novel intercellular signaling pathways and cohorts of extracellular ligands that act synergistically to affect developmental processes. While we employ this methodology towards the identification of gliogenic inducers, other biological questions within and outside the nervous system are almost unlimited. Most importantly, by combining ligand-receptor pairs that act on complimentary downstream target genes, this approach provides opportunities to identify groups of signaling factors whose synergistic effects may otherwise have been overlooked.

## METHODS

### Candidate Ligand Identification

Candidate ligands were identified using the NicheNet algorithm^41^ implemented in R version 3.6.2. Mouse data was derived from Zhang et al.^42^ and human from 3 single cell datasets of the developing fetal brain^43–45^. All input datasets were first count normalized using DESeq2^46^. Bulk fastq files were processed by trimming using Trimmomatic^47^, alignment using STAR^48^ to mm10 and hg19, respectively, and reads were summarized using featureCounts^49^. Human single cell data was downloaded in count matrix format and processed using the Seurat v3 pipeline^50^. Cell type populations were identified after using the “find markers” function and were assigned identities based on markers defined by the providing datasets. NicheNet inputs were as follows: sender cells—neuronal progenitors, immature inhibitory neurons, immature excitatory neurons, mature inhibitory neurons, mature excitatory neurons; receiver cells—radial glia, ventricular radial glia, outer radial glia, immature astrocytes. Each human single cell dataset was run through the NicheNet pipeline separately, and overlapping hits were ultimately consolidated into the final groups. Ligands with complementary predicted receptors and target genes were prioritized. A curated list of ∼500 previously identified immature astrocyte genes^51^ (**Supplemental Table 1**) was used as the gene set of interest to specify potential downstream targets of ligand-receptor signaling.

### Generation of cortical organoids

Human cortical organoids were formed from three human induced pluripotent stem cell (hiPSC) lines (8858.3, 2242.1 and 1363.1)^52^ following a previously published protocol^53^. All lines were genotyped by SNP-array to confirm genomic integrity and regularly screened for mycoplasma. iPSC colonies at 80-90% confluency were detached from culture plates using Accutase and were formed into 3D spheroids using AggreWell™ plates. Following 3D formation, spheroids were treated daily in neural induction media (DMEM/F12, KSR, NEAA, Glutamax, Pen/Strep, Beta-mercaptoethanol) supplemented with Dorsomorphin (Sigma, Cat. P5499-25MG, 5 µM) and SB-431542 (Selleck Chemicals, Cat. S1067, 10 µM) for 6 days. Following this treatment, organoids were treated daily with neural media supplemented with EGF and FGF2 for 10 days, and every other day for days 16-24. At day 25, organoids were treated every other day with neural media supplemented with BDNF and NT-3 to promote differentiation of progenitors. From day 43 onwards, organoids were fed every 3 days with neural media only.

### Organoid Ligand Exposures

Organoids were exposed to candidate ligands continually with media changes every other day for thirty-day periods between day 45-75, day 60-90, and day 90-120. In preliminary exposures ligands were added in two groups. Group 1 ligands were derived from mouse transcriptomic data and included: APP (R&D Systems, Cat. 3466-PI-010, 20ng/mL), APOE3 (Sigma-Aldrich, Cat. SRP4696-500UG, 30ng/mL), GAS6 (R&D Systems, Cat. 885-GSB-050, 80ng/mL), CALR (Abcam, Cat. ab91577, 15ng/mL), and IGF1 (Sigma-Aldrich, Cat. I3769-50UG, 100ng/mL). Group 2 ligands were derived from human transcriptomic data and included: TGFβ2 (R&D Systems, Cat. 302-B2-002, 5ng/mL), NLGN1 (Sino Biological, Cat. 11617-H08H-100, R&D Systems, Cat. AF4340, 50ng/mL), TSLP (Sino Biological, Cat. 16135-H08H, 20ng/mL), DKK1 (Sigma-Aldrich, Cat. SRP3258-10UG, R&D Systems, Cat. 5439-DK-010/CF, 20ng/mL), and BMP4 (Sigma-Aldrich, Cat. H4916-10UG, R&D Systems, Cat. 314-BP-010/CF, 10ng/mL). In successive exposures, ligands were added to cultures at these concentrations either in combination for “cocktail” exposures or individually.

### RNA-Sequencing Library Preparation

Following exposures, total RNA was extracted from organoids using the miRNeasy kit (Qiagen, Cat. 217084) under the protocols of the manufacturer (discarding microRNA fraction). The quality of the RNA was assessed by Bioanalyzer and only samples with RIN > 9.0 were used for library preparation. Bulk RNA-seq libraries were created with the NEB Next Ultra II kit using poly-A selection and sequenced to a depth of 40 million paired-end reads per sample. For the preliminary targeted RNA-seq experiment we generated a custom 100-gene panel using Qiaseq (Qiagen). This panel included 40 astrocyte-specific genes, 40 neuronal-specific genes, 10 housekeeping control genes (set by the manufacturer), and 10 reactive astrocyte genes^51,54^ **(Supplemental Table 2)**. These genes were selected by (a) their enrichment in astrocytes or neurons over other human cell type populations (including oligodendrocytes, endothelial cells, and myeloid cells), and (b) their baseline expression levels above the 50^th^ percentile. Together, these criteria make these genes easier to detect and highly cell type-specific.

### RNA-Sequencing Processing and Analysis

Targeted RNA-seq data was processed using the Qiagen GeneGlobe software and normalized using EdgeR. For bulk RNA-seq, fastq files were first trimmed using Trimmomatic and mapped using STAR aligner with the paired end option selected (hg19). We used FeatureCounts to assemble transcripts and generate raw count matrices. Following generation of raw count data, we used DESeq2 to normalize matrices and to determine differential gene expression statistics. To assess changes in astrocyte and neuronal gene signatures we assembled a list of 50 genes (25 astrocyte-specific and 25 neuronal-specific) **(Supplemental Table 3)** to use as a benchmark for comparing control and ligand-exposed conditions. Normalized count data for each gene was used to calculate fold changes as follows: (ligand exposure expression (TPM) / control condition expression (TPM)).

### Immunopanning organoid-derived and human fetal astrocytes

All human tissue samples were obtained in compliance with policies outlined by the Emory School of Medicine IRB office. Astrocytes were purified from human fetal tissue between 17-20 gestational weeks using a protocol outlined in ^51^. Tissue was chopped using a #10 blade and incubated in 7.5 U/ml papain at 34°C for 45 minutes before rinsing with a protease inhibitor solution (ovomucoid). After digestion, the tissue was triturated and then the resulting single-cell suspension was added to a series of plastic petri dish pre-coated with cell type specific antibodies and incubated for 10-15 minutes each at room temperature. Unbound cells were transferred to the subsequent petri dish while the dish with bound cells was rinsed with PBS to wash away loosely bound contaminating cell types. The antibodies used include anti-CD45 to harvest and deplete microglia/macrophages, anti–Thy1 to deplete neurons, and anti–CD49f to collect astrocytes. For RNA-seq, cells were directly scraped off the panning dish with Qiazol (Qiagen). For cell culture and *in vitro* experiments, astrocytes bound to the antibody-coated dishes were incubated in a trypsin solution at 37°C for 3–5 minutes and gently squirted off the plate. Cells were then spun and plated on poly-D-lysine-coated plastic coverslips in a Neurobasal/DMEM based serum-free medium. We replaced the media every other day for 12 days with or without ligand addition.

The same protocol described above was used for immunopanning organoids with minimal exceptions. These include smaller volume dissociations and spins, and omission of filtering steps that might reduce yield.

### Onset of gliogenesis experiments

Organoids from 3 previously validated hiPSC lines (8858.3, 2242.1 and 1363.1) underwent a total of 10 separate differentiations. Organoids were sampled from each differentiation at 10-day intervals from day 70 through day 110 (5 timepoints) and were administered 3 separate assays: quantitative real-time PCR (qPCR) for GFAP (protocol details below), immunohistochemistry IHC for GFAP (protocol details below), or immunopanned with anti-HepaCAM antibody. Criteria used to consider “successful” gliogenesis included a qPCR GFAP CT value < 28, greater than 5% GFAP+ cells as a percentage of total DAPI population, and immunopanning pulldown of at least 10,000 cells (∼5% yield).

### Immunocytochemistry

Cultured cells were fixed with 4% PFA for 10 minutes at room temperature, permeabilized and blocked with 10% donkey serum with 0.3% Triton-X100. Antibodies used were DAPI (in VECTASHEILD, Vector Laboratories, Cat. H-1500), GFAP (DAKO, Cat. Z0334, dilution 1:1500), and Ki67 (BD, Cat. b550609, dilution 1:50).

### EdU Exposure

Thymidine analogue EdU (Invitrogen, Cat. C10640) was added to every media change at a final concentration of 10 µM to label proliferating cells. EdU staining was performed according to manufacturer recommendations (Invitrogen, Cat. C10640) followed by immunocytochemistry. Total EdU-positive and DAPI-positive cell counts were calculated using the Keyence Hybrid Cell Count software.

### Immunohistochemistry

Organoids were fixed with 4% PFA for 30 minutes at room temperature, rinsed with PBS, and then equilibrated with 30% sucrose overnight. The following day, after organoids sunk to the bottom of the tube, they were embedded in OCT blocks and frozen for cryosectioning. 12µm sections were mounted on glass slides, permeabilized with triton-X100, and blocked with 10% donkey serum with 0.3% Triton-X100. Antibodies used included GFAP (DAKO, Cat. Z0334, dilution 1:1500). Coverslips were mounted with DAPI in VECTASHEILD (Vector Laboratories, Cat. H-1500).

### Quantitative real-time PCR

Total RNA was extracted from organoids using the miRNeasy kit (Qiagen Cat. 217084) under the protocols of the manufacturer (discarding microRNA fraction). The quality of the RNA was assessed by Bioanalyzer and only samples with RIN > 9.0 were used for library preparation. cDNA was synthesizing using a reverse transcriptase reaction with random hexamers and oligodT primers. We performed 40 cycles of amplification for all samples. The specificity and efficiency of all primers were first validated using gel electrophoresis and qRT-PCR with serial dilutions. The determination of each gene’s CT in qRT-PCR was performed in triplicate. When determining fold changes in gene expression across samples, the CT of each gene was normalized according to the CT of the housekeeping gene in the same sample. The primers used in this study include:

GFAP Forward: GAGAACCGGATCACCATTCC

GFAP Reverse: CCCAGTCTGGAGCAACCTAC

GAPDH Forward: AATCCCATCACCATCTTCCA

GAPDH Reverse: TGGACTCCACGACGTACTCA

### Astrocyte Morphology Quantification

Images of GFAP+ cells were traced and analyzed using the Fiji plug-in SNT. 20 cells from the control and candidate ligand exposed conditions were traced using SNT’s semi-automated tracing method. Primary branches originate at the nucleus. Secondary branches extend off primary branches, and tertiary branches extend off secondary branches. The SNT software quantified the total path number, primary, secondary, and tertiary path numbers, and path length for each cell.

### Phosphoproteomic array

The phosphorylation pathway profiling array was performed following all standard manufacturer’s instructions (RayBiotech AAH-PPP-1-2). Lysates were prepared from hCOs exposed to control or ligand supplemented conditions for 30 days. Provided lysis buffer and protease/phosphatase inhibitor cocktails were used to prevent further phosphorylation / dephosphorylation events during processing. Total protein concentrations for each sample were measured using BCA and normalized prior to loading membranes. All incubations and washes were performed according to manufacturer instructions with all antibody incubations performed at 4C overnight. Chemiluminescence was performed using a ChemiDoc imager. Image analysis was performed using Fiji to extract density information for each blot. Identical circle dimensions (area, size, shape) were used to measure signal densities across all arrays and summed signal density was calculated per spot according to manufacturer protocol. After determining signal densities for each spot, we performed background subtraction using the negative control spots followed by positive control normalization using positive control spots. This allowed us to reliably compare across multiple arrays analogous to housekeeping proteins on typical Western Blots. Signal fold expression was calculated using the following calculation (provided by manufacturer):

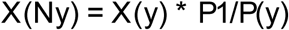

Where:

P1 = mean signal density of Positive Control spots on the reference array

P(y) = mean signal density of Positive Control spots on Array “y”

X(y) = mean signal density for spot “X” on Array “y”

X(Ny)= normalized signal intensity for spot “X” on Array “y”

## RESULTS

### NicheNet predicts ligand and receptor pairs that influence astrocyte development

In humans, neurogenesis temporally precedes astrogenesis. This switch in cell fate depends on both intrinsic and extrinsic signals that act in or on RG progenitors (**Figure 1A**). For an extrinsic signal to exert an astrogenic effect, it must meet three important criteria: first, it must be expressed by cells that are present prior to gliogenesis. Second, it must bind to cognate receptors that are expressed in astrocyte progenitors (*e*.*g*. RG) or early astrocytes. Finally, this ligand-receptor event must exert downstream changes that lead to the expression of astrocyte signature genes. To identify ligands that meet these criteria, we applied a computational discovery approach called NicheNet. This algorithm uses transcriptomic data as input (either bulk or single cell) to identify expressed ligands and their receptors in a tissue of interest. Furthermore, NicheNet uses existing knowledge of signaling networks to predict the effects of each ligand-receptor binding event on the downstream gene expression of a set of target genes (**Figure 1B**).

**Figure 1.**
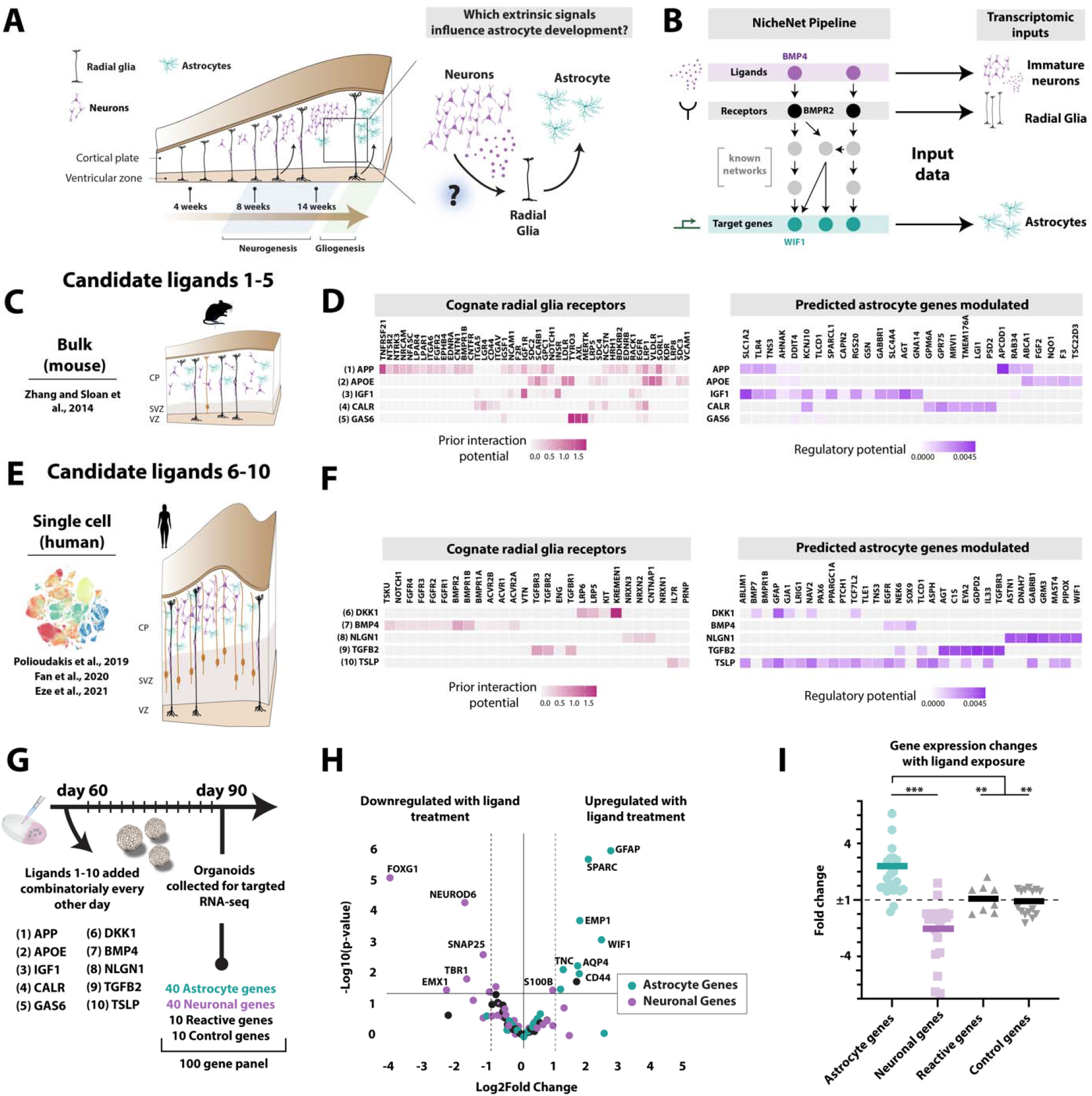
Computational identification of candidate gliogenic signals **A**. Neurogenesis precedes gliogenesis during human fetal brain development. **B**. NicheNet pipeline matches secreted ligands from a “sender cell” population with cognate receptors from a “receiver cell” population. The corresponding ligand-receptor pairs are then scored by their ability to regulate a defined gene set. In total these are the three inputs to the algorithm—sender cells, receiver cells, and a target gene set. **C**. Mouse RNA-seq data from Zhang and Sloan et al. was used as input data **D**. (Left) Interaction potential of top 5 ligands and their cognate receptors identified from the mouse data. (Right) Regulatory potential of the candidate ligands to specifically impact the target gene set. **E**. Human single cell RNA-seq data from Polloudakis et al., Fan et al., and Eze et al. was used as input data **F**. (Left) Interaction potential of top 5 ligands and their cognate receptors identified from the human single cell data. (Right) Regulatory potential of the candidate ligands to specifically impact the target gene set. **G**. Ligand exposure paradigm using a targeted RNA-seq readout. **H**. Volcano plot of targeted RNA-seq data from days 60-90 after ligand exposure to a cocktail of all 10 ligand candidates. All values are compared to control hCOs cultured in neural media with standard conditions. (n = 3 control and 3 experimental organoids sequenced separately) **I**. Quantification of gene expression changes from the targeted RNA-seq panel following ligand cocktail exposure. Mann-Whitney U test (*** p<.001, ** p<.01).

To identify a list of candidate ligands capable of modulating human astrocyte development, we applied NicheNet to both mouse and human datasets from the developing cortex (P3-7 for mouse and gestational week 16-19 for human). The rationale for beginning with mouse data is that it provided a source of deeply sequenced purified cell type-specific inputs, whereas human datasets are largely restricted to low-depth single cell information. Therefore, we hypothesized that using both types of data as separate inputs would serve as a valuable screen for effectors of astrocyte development. For this study, we focused on neuronal populations as our “sender cells” (ligand sources), although it is well-established that other CNS populations can also contribute to astrocyte development. We defined radial glia (both ventricular and outer) and immature astrocytes as “receiver cells”. NicheNet’s third and final input is a target gene-set that can be used to benchmark the regulatory potential of each ligand-receptor pair. To define this target gene-set, we identified a list of 500 astrocyte-specific genes^51^ spanning both immature and mature developmental stages **(Supplemental Table 1)**. From these inputs derived from the mouse dataset, we generated a list of ligand-receptor pairs that (1) are expressed in relevant cell types, (2) exhibit binding interaction, and (3) are predicted to act upstream of astrocyte-specific target genes. From this list, we narrowed to a group of 5 ligands (1-5; APP, APOE, IGF1, CALB, GAS6) by focusing on candidates exhibiting complementary receptor binding and activation signatures of astrocyte genes (**Figure 1C-D**). We specifically selected ligands that act on separate signaling pathways and promote distinct astrocyte signature genes because they are more likely to work synergistically to drive astrocyte development.

For our human analysis, we explored single cell data from three separate studies of developing human fetal brain tissue (**Figure 1E**). We assigned all cell IDs to a specific cell type identity based on classic cell type markers (cite). For our sender cell population, we included all immature and mature neuronal subtypes (excitatory, inhibitory, and intermediate progenitors). For receiver cells, we included ventricular radial glia, outer radial glia, and immature astrocytes. Our target gene-set of interest remained our 500 gene human astrocyte-specific signature. This analysis yielded a separate set of candidate ligands (6-10; DKK1, BMP4, NLGN1, TGFβ2, TSLP), again selected for their complementary receptor binding and activation signatures of astrocyte genes (**Figure 1F**). Of note, both BMP and TGFβ signaling have been well-implicated as modulators of astrocyte development^31–33^, but these molecules had not previously been investigated for their potential synergistic contributions.

As a preliminary screen of these candidate ligands, we exposed human cortical organoids (hCOs) across three hiPSC lines to a cocktail of all 10 ligands. Bioactive ligand concentrations were extrapolated from the literature and empirically tested *in vitro* to ensure no toxicity issues **(Supplemental Table 4)**. The hCO ligand exposures occurred over a 30-day period spanning days 60-90 *in vitro*, prior to the onset of gliogenesis. Ligands were added to the media every other day to maintain stable levels. To effectively readout whether this ligand cocktail influenced the balance between neuronal and glial commitment, we designed a custom targeted RNA-seq panel (Qiagen) containing 40 astrocyte genes, 40 neuronal genes, 10 reactive astrocyte genes, and 10 control housekeeping genes (**Figure 1G, Supplemental Table 2**). Using a large gene panel helps ensure that our interpretation is not biased by the specific induction of a small subset of 2-3 marker genes that could be prone to individual bias by a single ligand (i.e. GFAP). After a 30-day exposure to the ligand cocktail, targeted sequencing revealed a significant upregulation of astrocyte genes (Mann Whitney U, p < .0001) and a concomitant downregulation of neuronal genes (Mann Whitney U, p < .0001). No control (p = .765) or reactive genes (p = .881) were significantly changed upon ligand exposure (**Figure 1H-I)**.

### Only human identified gliogenic ligands influence astrocyte development

We next wondered whether the ligands identified through the mouse data (1-5) or human single cell datasets (6-10) were specifically driving the expression changes that we observed in our larger screen. Therefore, we split our ligand cocktails into these two groups and performed identical 30-day exposures of hCOs. Again, using an astrocyte and neuronal signature gene-set (**Supplemental Table 3**) as the readout, we observed that only the human-identified ligands (6-10) induced a robust response (Mann Whitney U, p = .002) (**Figure 2A-C**), whereas there was no change in the presence of the mouse-identified ligands (Mann Whitney U, p = .781). The effect of the group 2 (human-identified) ligands alone was also comparable to the significant effects seen with all 10 ligands added together, suggesting that the group 1 (mouse-identified) ligands were unlikely to be modulating the effects. Therefore, we proceeded with ligands 6-10 and refer to these as our human-derived ligand cocktail for all subsequent experiments.

**Figure 2.**
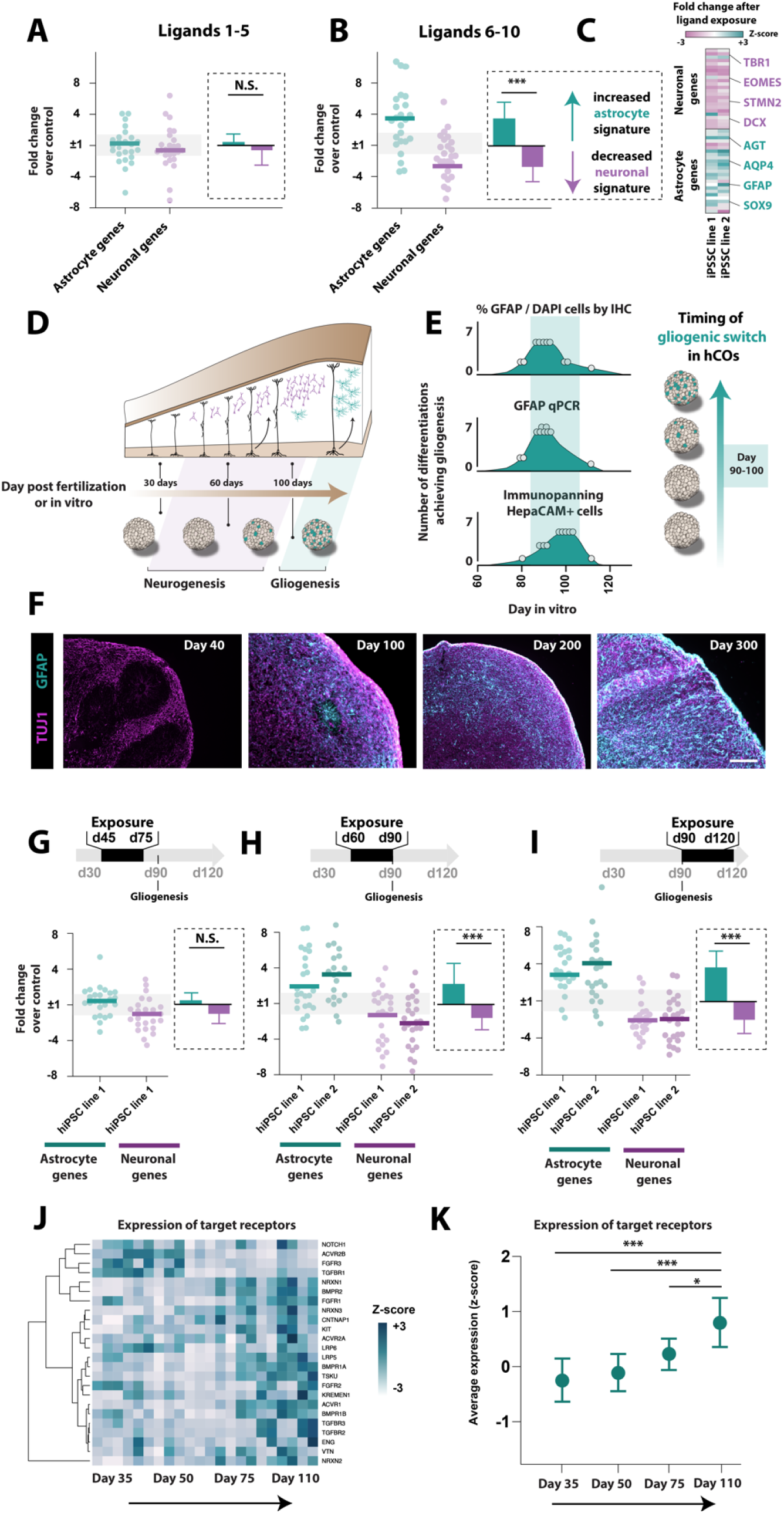
Ligand activity depends on hCO developmental stage. **A**. Following a 30-day exposure in hCOs, mouse-identified ligands 1-5 do not significantly modulate astrocyte or neuronal gene signatures (N.S. p = .781, Mann Whitney U) (n = 8 control organoids, 12 ligand-exposed organoids). **B**. Human-identified ligands 6-10 significantly increase expression of astrocyte gene signatures and decrease expression of neuronal genes (p = .002, Mann Whitney U). **C**. Heatmap of neuronal and astrocyte gene expression following ligand exposure. Classic signature genes for each population are highlighted. **D**. Schematic of proposed timeline of the gliogenic switch in organoid cultures. **E**. Variability of gliogenic switch in hCOs. 10 separate hCO differentiations (n = 5 hiPSC lines) were assayed for (1) % GFAP+ cells by IHC, (2) GFAP mRNA by qPCR, and (3) number of cells bound to HepaCAM+ immunopanning plate at 10-day intervals from days 70-110 in culture. Thresholds were set for each assay to determine that gliogenesis had begun. These include >5% GFAP/DAPI+ cells for IHC (top panel), a CT cutoff <30 for GFAP qPCR (middle panel), and >10,000 HepaCAM immunopanned astrocyte per organoid (lower panel). **F**. Representative GFAP and TUJ1 staining of day 40, day 100, day 200, and day 300 hCOs. Scale bar =100 µm, **G-I**. Result of ligand exposures (6-10) at different stages of hCO culture. Readouts are fold change of astrocyte and neuronal gene signatures compared to control hCOs using RNA-seq. (N.S. p = .643, ***p<.0001, Mann Whitney U) (n = 6-8 control organoids and 6-12 experimental organoids). **J-K**. Expression of target receptors for NicheNet-predicted ligands 6-10 throughout hCO development (***p<.0001, ***p<.0001, *p = .013, Mann Whitney U).

### The gliogenic switch occurs reproducibly around day 90 in hCOs

To determine if there is a temporal window during which candidate ligands most potently influence astrocyte development, we first needed to precisely define the onset of gliogenesis within hCOs. There are numerous metrics that have been used to define the initiation of astrocyte formation, each with their own caveats and advantages. Therefore, we chose to use three separate assays to be as comprehensive as possible in our definition of gliogenesis. These include (1) immunohistochemistry to quantify the abundance of GFAP+ cells, (2) qPCR to quantify total GFAP mRNA within hCOs, and (3) immunopanning to pulldown HepaCAM^**+**^ astrocytes. For each metric, we set thresholds based upon values observed in human fetal tissue at gestational week 17 when gliogenesis is initiated (see Methods). Next, we generated 10 separate differentiations of hCOs across 4 hiPSC lines and assayed for each of the above outcomes at days 70, 80, 90, 100, and 110. From these assays, we created a temporal map of the onset of gliogenesis based on outcomes from each separate criterion. Remarkably, the onset of gliogenesis was reproducible and consistent across lines, differentiations, and outcome metrics at a time window between day 90-100 of hCO culture (IHC for GFAP^**+**^ cells: 92 ± 9 days, qPCR for GFAP: 99 ± 6 days, immunopanning for HepaCAM^**+**^ cells: 96 ± 8 days) (**Figure 2D-F**).

### Ligand exposures affect astrocyte development before and after the gliogenic switch

Based on this timeline of astrogenesis, we wondered whether the ligand cocktail would exhibit differential effects when exposed to hCOs at timepoints far preceding (day 45-75), immediately before (day 60-90), and immediately after the onset of gliogenesis (day 90-120). Selection of these timepoints allowed us to both assay the ligands’ developmental effects and test the temporal receptivity of RG to these signals. During exposures that lasted from day 45-75, we found no significant difference (Mann Whitney U, p = .643) between astrocyte and neuronal gene expression. However, at the day 60-90 and day 90-120 exposures, we observed a significant increase in astrocyte genes and concomitant decrease in neuronal gene expression (Mann Whitney U, p ≤ .0001 at each timepoint) (**Figure 2G-I**).

### The cognate receptors of the ligands are developmentally regulated

Given the susceptibility of cells to respond to the ligand cocktail only at timepoints before and after the gliogenic switch, we predicted that the expression patterns of the cognate receptors to these ligands might correlate with developmental stages. To test this hypothesis, we performed bulk RNA-seq of whole hCOs to analyze receptor expression at various developmental timepoints (day 35, day 50, day 75, day 110). We found that the majority of the predicted ligand-binding receptors increase in expression as hCOs approach gliogenic timepoints (**Figure 2J-K**). Thus, the lack of significant changes in astrocyte and neuronal gene expression following ligand exposures from day 45-75 could be the result of low expression of receptors on radial glia at these timepoints. These data further confirm that the day 60-90 and day 90-120 exposures fall within a key period for astrocyte development.

### Ligands work synergistically, but not individually, to influence astrocyte development

Given our findings that our ligand cocktail supports astrocyte development at exposures from day 60-90 and day 90-120, we next sought to compare the impact of synergistic ligand administration versus each individual ligand on astrocyte development. We performed ligand exposures from day 60-90 (**Figure 3A**) or day 90-120 (**Figure 3D**), either adding our candidate ligand cocktail, or adding each ligand individually to hCOs. Of the ligands added individually, only the addition of BMP4 resulted in a significant increase in astrocyte gene signatures and concomitant decrease in neuronal gene signatures. However, at all timepoints, we observed that the degree of astrocyte signature induction and neuronal signature depletion was most significant with all 5 ligands combined (**Figure 3B-C, 3E-F**).

**Figure 3.**
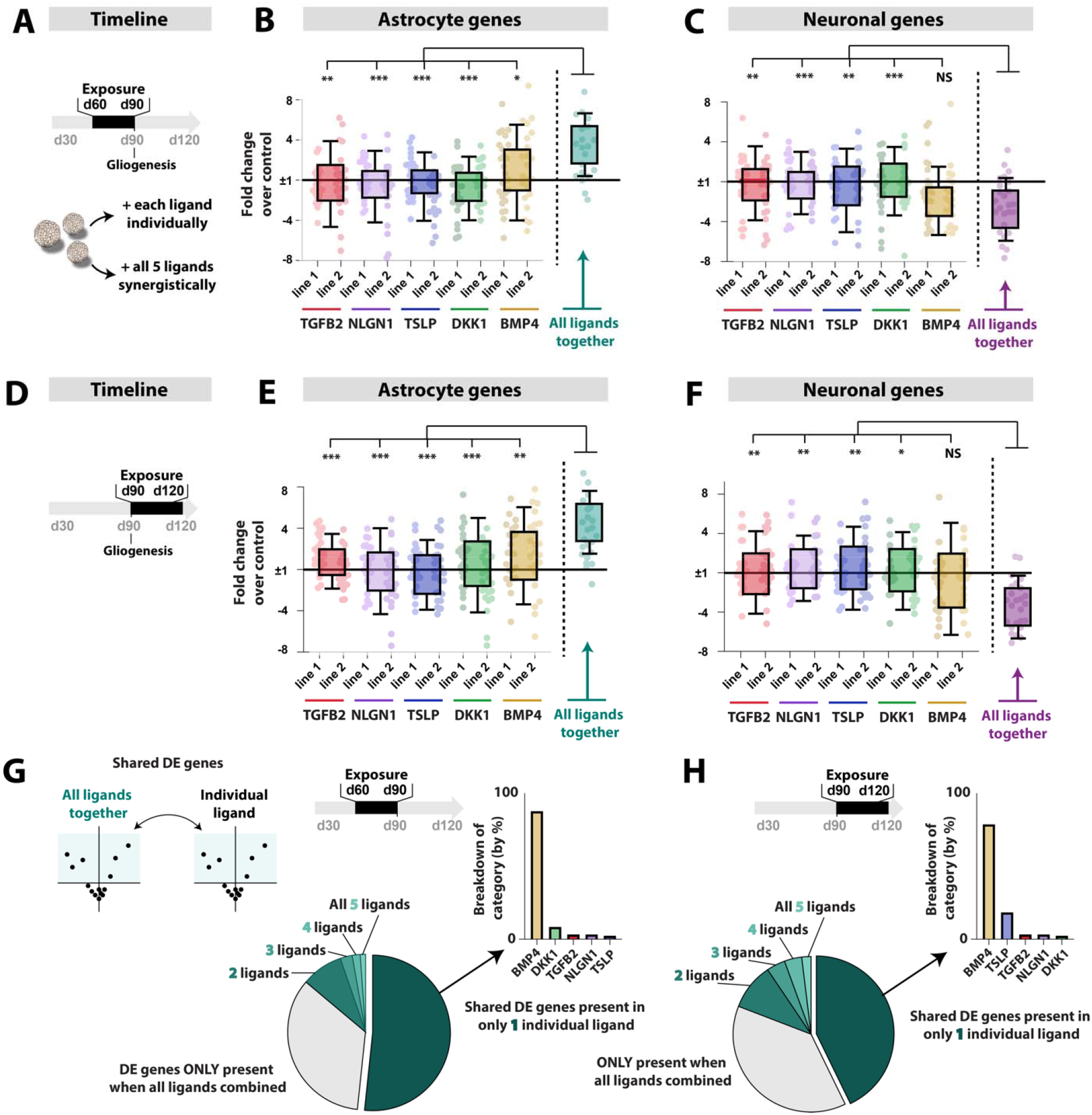
Synergistic and individual contributions of ligands 6-10. **A**. Schematic of individual ligand exposure paradigm. Ligands were added every other day between day 60-90 either individually or combined together in a 5-ligand cocktail (n = 8 control, n = 4 ligand exposed organoids). **B**. Astrocyte gene signatures in the presence of each individual ligand or the full cocktail (*** p<.0001, **p<.001, *p<.01, Mann Whitney U) **C**. Neuronal gene signatures in the presence of each individual ligand or the full cocktail. **D**. Timeline of ligand administration for later timepoint exposure. Ligands were added every other day between days 90-120 either individually or combined together in a 5-ligand cocktail (n = 8 control, n = 4 ligand exposed organoids). **E**. Astrocyte gene signatures at day 120 in the presence of each individual ligand or the full cocktail. **F**. Neuronal gene signatures at day 120 in the presence of each individual ligand or the full cocktail. **G**. Differential expressed genes (both up and down, as defined by p-adjusted < .01 and log2FC > 2) were identified in hCOs treated with all 5 ligands and each ligand separately. Pie chart illustrates the percent of DEGs that overlap with no individual ligand conditions, only 1 condition, or 2-5 conditions. Of the DEGs present in only 1 individual ligand condition, the distribution of those genes are subset in the bar charts.

We next wondered how transcriptomic changes induced by the ligand cocktail could be explained by gene changes produced by each ligand separately. We first identified all differentially expressed genes (up or down) in the presence of the ligand cocktail (n = 817 and 542 genes at day 60-90 and 90-120, respectively) and asked which of these genes were also dysregulated in 1, 2, 3, 4, or all 5 individual ligand conditions. Regardless of timepoint administration, we found that the majority of cocktail-induced genes changes were also differentially expressed in at least one individual ligand condition (78%). Of these, the vast majority were perturbed in only one single ligand (57%), compared with 21% in 2 or more ligands. Interestingly, another 22% of genes (∼150) were only dysregulated when all 5 ligands were added together (**Figure 3G-H)**.

Finally, we wondered if certain individual ligands were contributing more than others to the cocktail-induced changes. We specifically subset those genes that were both differentially expressed in the ligand cocktail condition and only one single ligand exposure condition. Of these transcriptomic changes, BMP4 was the predominant source (**Figure 3G-H)**. Importantly, this accounted for only ∼35-40% of the overall gene changes observed in the ligand cocktail.

### Candidate ligands also impact astrocyte development in fetal human samples

With evidence of transcriptomic changes in hCO-derived astrocyte populations, we next aimed to benchmark the hCO model against primary human astrocytes. This comparison offers the added benefit of testing whether ligand perfusion into a 3D structure might impact their potency. We purified CD49f+ astrocyte populations from human fetal brain tissue collected between 17-20 gestational weeks using immunopanning. Following purification, astrocytes were cultured in monolayer for 10-12 days with ligand exposures occurring every other day to maintain stable levels (**Figure 4A)**. Following ligand exposure we performed RNA-seq of purified cells and again observed a striking induction of astrocyte genes and downregulation of neuronal genes (Mann Whitney U, p <.001) (**Figure 4B)**.

**Figure 4.**
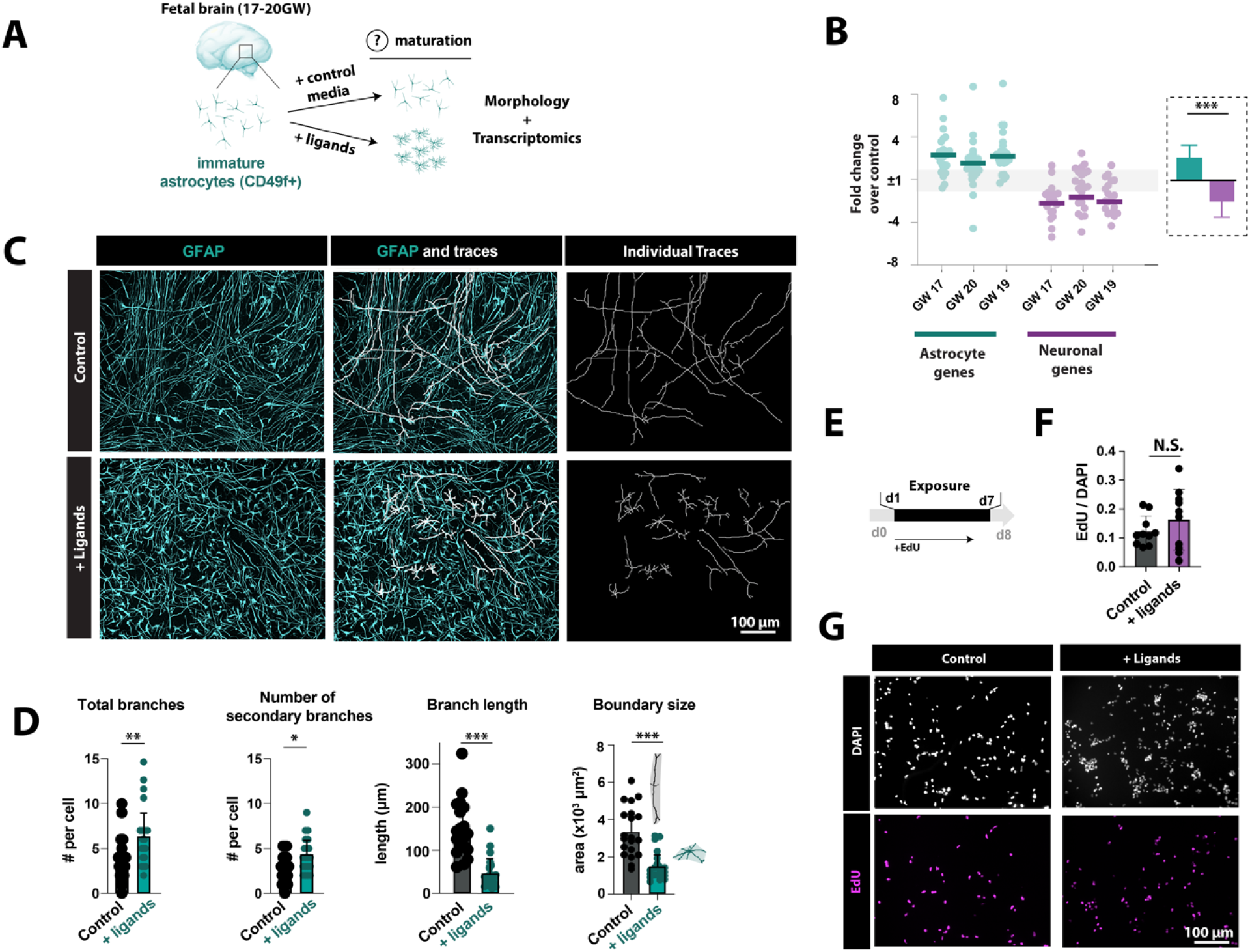
Impact of candidate ligand exposures on fetal astrocytes. **A**. GW 17-20 cortices were immunopanned for CD49f+ immature astrocytes, which were cultured for 10 days in the presence or absence of the ligand cocktail. **B**. Astrocyte and neuronal gene signatures assessed by RNA-seq of CD490f+ cells. Fold change represents expression in ligand conditions vs control media (p < .001, Mann Whitney U). **C**. GFAP+ cell process traces from ligand-exposed and control CD49f+ fetal cells. Scale bar = 50 µm. **D**. GFAP+ cell process quantification. Primary branches extend from the nucleus. Secondary branches extend from primary branches. Boundary size is the area (x 10^3^ µm^2^) of the image field that one cell occupies. (*p<.01, **p<.001, *** p<.0001, Mann Whitney U). **E**. Timeline of EdU exposure. Fetal astrocytes were cultured for 8 days after purification with ligand exposure from days 1-7. EdU added at day 1 until duration of experiment. **F**. No significant change in percent of EdU+ nuclei between control and ligand-exposed cells. **G**. Representative images of DAPI and EdU+ cells after 8 days in culture.

### Synergistic ligand exposure drives mature astrocyte morphology in purified fetal astrocytes

We next aimed to understand the effects of the ligand cocktail on astrocyte morphology. Investigating morphology allowed us to quantify the effects of the ligands on physical astrocyte structure, which can be a useful indicator of astrocyte maturation. Radial glia and immature astrocytes typically exhibit a more bipolar and elongated morphology, while mature astrocytes have a more branched, star-shaped morphology^55,56^. We cultured purified CD49f+ fetal cells (17-20 GW) for 10-12 days in the presence or absence of our ligand cocktail. We then fixed these ligand-exposed fetal astrocyte cultures and used immunohistochemistry to visualize the morphology of the major branches of each GFAP+ cell (**Figure 4C**). We used semi-automated tracing of astrocyte processes to quantify the number, length, complexity, and boundary area of astrocyte branching. Using these outputs, we found a significantly increased number of total branches (Mann-Whitney U test, p = .004), increased number of secondary branches (Mann-Whitney U test, p = .02), decreased branch length (Mann-Whitney U test, p = .001), and decreased boundary size (Mann-Whitney U test, p = .027) compared to control fetal astrocyte cultures (**Figure 4C-D**).

### Synergistic ligand exposure does not affect fetal astrocyte proliferation

Given the profound effect of the ligand exposure on inducing astrocyte gene expression, we next wondered whether these ligands induced proliferation of fetal astrocytes. To test this, we purified CD49f+ fetal cells (17-20 GW) and cultured with the thymidine analogue EdU for 8 days in the presence or absence of our ligand cocktail (**Figure 4E**). We then fixed these ligand-exposed fetal astrocyte cultures and used immunohistochemistry to visualize and quantify the percentage of proliferating cells (EdU+/DAPI+). We found no significant difference between the control and ligand exposed cells, suggesting the ligands do not act directly on astrocyte proliferation (**Figure 4F-G**).

### Extrinsic cocktail of gliogenic ligands converge on modulating AKT/mTOR signaling

Since the gliogenic cocktail was capable of modulating astrocyte transcriptional programs, morphology, and proliferation, we next wondered which downstream pathways might mediate these changes. To supplement our transcriptional analytics, we pursued an approach to investigate the phospho-proteome of key signal transduction pathways in response to ligand exposure. Our hypothesis was that synergistic ligand cooperativity might converge on activation of specific pathway(s) that are essential for driving astrogenesis and maturation. Furthermore, since we selected this ligand cocktail by their divergent receptor repertoires, we predicted that their synergistic effects may act broadly on multiple pathways. To test this, we used a human phosphorylation pathway profiling array (RayBioTech) to simultaneously measure 55 different protein phosphorylation events across five major signal transduction pathways (BMP, AKT/mTOR, JAK/STAT, TGFβ, and NF-κB) (**Figure 5A-E)**.

**Figure 5.**
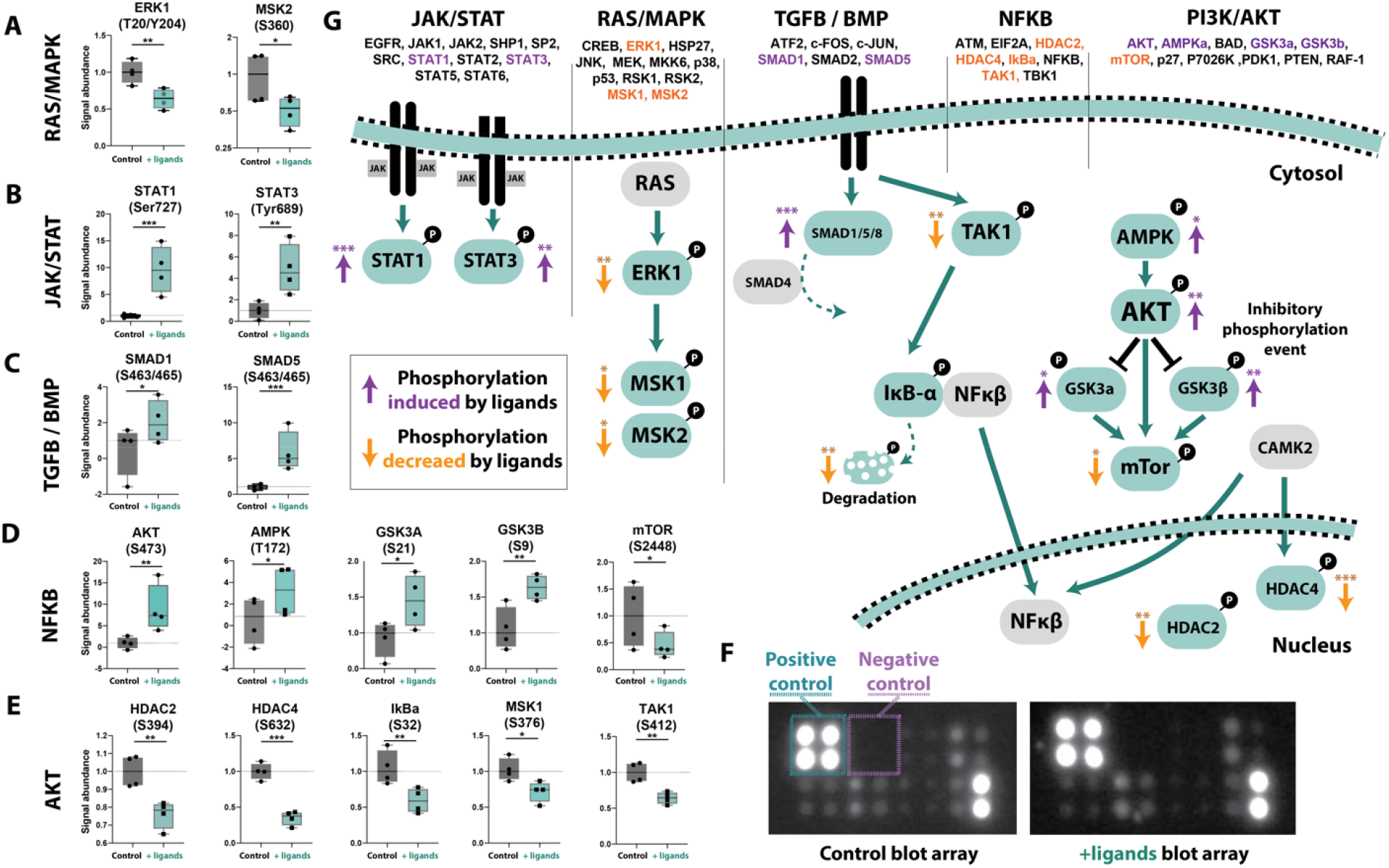
Phosphoproteomic changes in ligand-treated hCOs. **A-E**. Protein abundances between control and ligand cocktail treated hCOs. All values were normalized to control signal. Specific probed phosphorylation sites for each protein are listed under the protein name. Only significant changes are displayed in each of the five assayed pathways. (*p<.01, **p<.001, *** p<.0001, Mann Whitney U). **F**. Raw images of control and ligand-treated blot arrays, which include positive and negative controls to scale values on each individual array. **G**. Visual schematic of major signaling pathway changes in the presence of gliogenic ligand cocktail. Phosphorylation events induced by ligands are shown in purple and those decreased by ligand exposure are illustrated by orange arrows. All significantly changed proteins are shown in teal. Those in grey did not exhibit changes upon ligand administration.

We exposed hCOs to the gliogenic cocktail for 30 days (day 60-90) and then harvested total protein before normalizing inputs and performing the dot blot arrays (**Figure 5F**). Of the 55 probed protein phosphorylation events (n = 2 hiPSC lines, 4 replicates), we observed significant changes across 16 separate phosphorylated proteins belonging to all 5 signaling pathways (**Figure 5A-E)**. Some of these phosphorylation events were predicted direct downstream consequences from ligand activation (SMAD1 from BMP4, STAT3 from TGFβ2), while numerous others were likely the result of synergistic pathway activation. Most notable was the regulation of the AMPK/mTOR signaling pathway (**Figure 5D-E**). Of the 14 phosphorylation events in this array, 6 were significantly dysregulated upon ligand exposure. This included a 2.8-fold increase in AMPK phosphorylation (p = .041), 1.4-fold increase in GSK3a (inactivating phosphorylation event, p = .038), 1.7-fold increase in GSK3b (inactivating phosphorylation event, p = .031), and 2.2-fold decrease in mTor phosphorylation (p = .044). Inactivation of mTor phosphorylation in ligand-exposed hCOs is consistent with accelerated maturation and decreased proliferation of astrocyte progenitors and could explain a synergistic consequence of the ligand cocktail. In addition to downregulated mTor activity, we also observed a significant decrease in two histone deacetylase enzymes, HDACs 2 and 4 (p = .008, .002, respectively) that act downstream of AKT/mTOR signaling. Taken together, we found that exposure to the gliogenic ligand cocktail leads to activity changes across multiple signaling pathways, and especially decreased mTOR/AKT signaling (**Figure 5G**).

## DISCUSSION

Cell fate decisions during organogenesis are driven by both intrinsic and extrinsic mechanisms. While many new genomic technologies are improving our ability to disentangle intrinsic drivers of cell lineage commitment, our capacity to identify novel extrinsic signals has not grown as rapidly. This largely results from the fact that there are thousands of putative secreted molecules throughout development, each of which exhibit their own temporal dynamics. Furthermore, disentangling how these ligands act cooperatively to affect changes in recipient cells has remained an ongoing challenge. Here, we use computational tools to predict a group of synergistically-acting ligands on astrocyte development. Specifically, we show that TGFβ2, NLGN1, TSLP, DKK1, and BMP4 can work cooperatively to influence astrocyte development. Interestingly, with the exception of BMP4, each of these ligands exhibit minimal effects on their own. Their combinatorial influence is far greater than the sum of each individual ligand.

### Mouse vs human identified ligands

When we began our search for gliogenic ligands, we started by using both mouse and human datasets. However, none of our top 5 ligands identified from the mouse data demonstrated an empirical effect on human astrocyte development. We hypothesize there are several possibilities for this outcome. First, it is possible that species differences in the expression profiles of neuronal and radial glial populations between mouse and human are divergent enough to obfuscate the NicheNet output^51,57–60^. Second, the mouse data had poorer cell type resolution than the human single cell data, and thus it is possible that the relevant cell type populations that are responding to extracellular ligands were missed or diluted in the mouse data. Finally, it is possible that the time points used in the mouse collected data (P6-7) were too late, given that the onset of gliogenesis in mice is closer to E18-P3^61^. In contrast, the human datasets we used were selected because they included data from fetal development specifically around the onset of gliogenesis. Altogether, we cannot easily distinguish between these or other possibilities for the inability of the mouse data to generate viable candidate ligands but believe that other rodent datasets could be valuable for predicting human ligand-receptor interactions if matched appropriately.

### Synergistic mechanism of candidate ligands

Our ligand cocktail contains molecules with a disparate range of attributes. TGFβ2 and BMP4 are both members of the TGFβ superfamily and mediate their transcriptional effects via both canonical (SMAD) and non-canonical (ERK/JNK) signaling pathways^62^. Interestingly, TGFβ2 and BMP4 can act both synergistically and antagonistically in various settings and tissues^63^. DKK1 is a negative regulator of Wnt signaling^64^, while TSLP is a pleiotropic cytokine that has been traditionally implicated in T-cell maturation and proinflammatory immune responses^65^. Finally, NLGN1 is a postsynaptic cell adhesion molecule important for neuronal spinogenesis, synaptic formation, and even astrocyte morphogenesis^66^. While it is difficult to postulate the exact mechanism by which these 5 ligands act cooperatively to promote astrocyte development, our protein phosphorylation assays suggest a potential convergence on AKT/AMPK/mTOR signaling. TGFβ2, BMP4, DKK1 and even NLGN1 have each been linked to mTOR activity via both direct and indirect regulation, suggesting that this may be an important mechanism by which this gliogenic cocktail influences cell fate commitment. Alternatively, it is also possible that each ligand exerts an individual effect on separate target genes, which then converge to transcriptionally drive a larger gliogenic effect. This could be mediated by mechanisms in which regulation of chromatin modifiers like HDAC2 and 4 endow a more permissive genomic landscape for alternative signaling pathways like BMP/TGFβ to exert an effect.

One of the ligands in our cocktail, BMP4, has been particularly well-studied and implicated in its role as an inducer of astrocyte maturation^31–33,35^. While we did observe a significant transcriptomic effect on astrocyte development when BMP4 was administered alone in our experiments, these changes were greatly eclipsed by the 5-ligand cocktail. Interestingly, some genes like GFAP, whose promoter is directly activated via BMP4-SMAD dependent signaling^37,39^, showed more similar activation levels in the ligand cocktail vs BMP4-only conditions. These data suggest to us that the greater impact of the synergistic ligands may be their ability to activate a wider swath of astrocyte genes rather than cooperatively induce higher expression of a smaller subset of astrocyte-specific genes.

### Consistency of gliogenesis in *in vitro* cultures

One of the more surprising findings from our study was the reproducibility of the timing of gliogenesis within hCOs across different hiPSC lines, sex, and differentiations. Given that heterogeneity has remained an important caveat of organoid cultures, it is intriguing that the timing of the cell fate change from neurons to glia is so preserved within this *in vitro* system. This is also consistent with observations from other 2D and 3D studies of neural differentiation^52,67,68^. Together, this supports previous data that the timing of gliogenesis is in part a result of intrinsic mechanisms that act as a *de facto* clock. However, from the extrinsic perspective, it suggests that the signals required to induce this fate change are present within this *in vitro* system. Thus, while other factors like hormonal regulation from the vasculature or immune modulation from microglia may have the capacity to modulate the timing of gliogenesis, they are not necessary for tight regulation of this process.

### Where do gliogenic signals arise?

In this study we specifically focused on neuronally-secreted ligands in the developing brain. However, the early CNS contains many other cell populations—microglia, endothelial cells, pericytes, and even other progenitors that could act via autocrine functions. This approach of matching secreted ligands with expressed receptors and their target genes could easily be applied to any and all of these cellular sources. In our model, hCOs are largely devoid of these non-ectodermal populations yet still undergo gliogenesis on a predictable timescale. For this reason, we hypothesized that at least some external cues must be neuronal in origin. One additional platform that may accelerate the discovery of extrinsic signaling cues that drive developmental paradigms is the use of spatial transcriptomic datasets. In our study we lacked spatial information about where these ligands were expressed within the brain microenvironment or even major regions, but new spatial datasets could allow for more sophisticated computational predictions about the physical juxtaposition of specific ligands and cognate receptors in cell types of interest.

### Conclusion

By applying a computational methodology to single cell and bulk developmental brain RNA-seq data, we predicted, tested, and validated a set of ligands that influence astrocyte development with both human organoids and primary fetal astrocytes. These ligands-TGFβ2, NLGN1, TSLP, DKK1, and BMP4-act synergistically to induce the expression of astrocyte genes far exceeding their cumulative individual capacity. In addition to their transcriptional effects, these ligands promote aspects of astrocyte maturation including morphological complexity and diminished proliferative ability. Altogether, this approach of mining existing datasets to identify and test extrinsic signals that drive cell fate changes is ripe for discovery. There are hundreds of possible ligand-receptor pairs and many additional potential combinations of these signals. Our study demonstrates the feasibility and efficacy of prioritizing these combinations to identify cohorts of ligands with important biological effects.

## Acknowledgements

We would like to thank the Sloan laboratory for helpful discussions and Andy Erwood for his contributions to early ligand identification. We would also like to thank Joseph Lee for his support with AWS management and usage. This study was supported by National Institute of Health grants (to SAS): R01MH125956 and R01NS123562, and Brain and Behavior NARSAD Young Investigator Award. AV is supported by the Barry Goldwater scholarship and the Emory IMSD program.

## Authors’ contributions

AJV and SAS designed all experiments. AJV and SNL performed ligand exposures. AJV, AK, and AS performed hiPSC cultures, organoid formation, and maintenance. AJV, MMS, EH, and CS assisted with fetal tissue preparations and experimental procedures. Bioinformatic processing was performed by AJV. AJV and SAS wrote the manuscript with input from all contributing authors.

