## Supplemental Table 1 for "Computational Identification of Ligand-Receptor Pairs that Drive Human Astrocyte Development"

List of all astrocyte-enriched genes used as the “geneset” in the NicheNet pipeline. These genes are then separated by their enrichment in either immature or mature human astrocytes from Zhang and Sloan et al. 2016.

| **Human astrocyte gene signature** | **Immature** | **Mature** |
| --- | --- | --- |
| A2ML1 |  | yes |
| ABCD2 |  | yes |
| ACBD7 |  | yes |
| ACOT11 |  | yes |
| ACOX2 |  | yes |
| ACSBG1 |  | yes |
| ACSL6 |  | yes |
| ACSS1 |  | yes |
| ACSS3 |  | yes |
| ADCYAP1R1 |  | yes |
| ADORA2B |  | yes |
| AGT |  | yes |
| AGXT2L1 |  | yes |
| AIFM3 |  | yes |
| AIMP2 | yes |  |
| AK4 |  | yes |
| ALDH1L1 |  | yes |
| ALDH4A1 |  | yes |
| ALDOC |  | yes |
| ANGPTL1 | yes |  |
| ANKFN1 | yes |  |
| ANKRD45 |  | yes |
| ANP32AP1 | yes |  |
| APOE |  | yes |
| AQP1 |  | yes |
| AQP4 |  | yes |
| ARC | yes |  |
| ARHGAP5-AS1 |  | yes |
| ARHGEF26 |  | yes |
| ARHGEF26-AS1 |  | yes |
| ASPM | yes |  |
| ATP13A4 |  | yes |
| ATP1A2 |  | yes |
| ATP1B2 |  | yes |
| AURKB | yes |  |
| B4GALNT4 | yes |  |
| BBOX1 |  | yes |
| BHMT2 |  | yes |
| BIRC5 | yes |  |
| BMPR1B |  | yes |
| BOC | yes |  |
| BUB1 | yes |  |
| BUB1B | yes |  |
| C11orf70 | yes |  |
| C11orf93 |  | yes |
| C14orf162 | yes |  |
| C15orf42 | yes |  |
| C16orf74 | yes |  |
| C16orf89 |  | yes |
| C17orf90 | yes |  |
| C19orf6 | yes |  |
| C1orf158 | yes |  |
| C4orf48 | yes |  |
| C7orf55 |  | yes |
| CA12 | yes |  |
| CABLES1 |  | yes |
| CACHD1 |  | yes |
| CACNB1 | yes |  |
| CASC5 | yes |  |
| CBS | yes |  |
| CCDC80 | yes |  |
| CCDC88C | yes |  |
| CCL27 | yes |  |
| CCNB2 | yes |  |
| CD24 | yes |  |
| CD38 |  | yes |
| CDC20 | yes |  |
| CDC45 | yes |  |
| CDC6 | yes |  |
| CDCA3 | yes |  |
| CDCA5 | yes |  |
| CDCA7 | yes |  |
| CDCA8 | yes |  |
| CDH4 | yes |  |
| CDON | yes |  |
| CELSR1 | yes |  |
| CENPE | yes |  |
| CENPF | yes |  |
| CENPM | yes |  |
| CERS1 |  | yes |
| CHI3L1 |  | yes |
| CHPF | yes |  |
| CHRDL1 |  | yes |
| CITED1 | yes |  |
| CKAP2L | yes |  |
| CKB | yes |  |
| CLDN10 |  | yes |
| CLU |  | yes |
| CMTM1 | yes |  |
| CNGA3 | yes |  |
| CNTFR | yes |  |
| CNTNAP3 |  | yes |
| COL4A6 | yes |  |
| COL5A3 |  | yes |
| COL6A1 | yes |  |
| CPE |  | yes |
| CRHR2 | yes |  |
| CST3 |  | yes |
| CTNNA2 |  | yes |
| CXCR7 |  | yes |
| CYP4F11 |  | yes |
| DBN1 | yes |  |
| DCHS2 |  | yes |
| DCT | yes |  |
| DDAH1 |  | yes |
| DDIT4L |  | yes |
| DIAPH3 | yes |  |
| DIO2 |  | yes |
| DLGAP5 | yes |  |
| DMRTA2 | yes |  |
| DNAH7 |  | yes |
| DOK5 | yes |  |
| DPYSL5 | yes |  |
| DTL | yes |  |
| DTX1 |  | yes |
| DTYMK | yes |  |
| DUS3L | yes |  |
| DUSP8 | yes |  |
| E2F7 | yes |  |
| EDNRB |  | yes |
| EFEMP1 |  | yes |
| EFHC2 |  | yes |
| EHHADH |  | yes |
| EIF3CL | yes |  |
| ELOVL2 |  | yes |
| EMX2 |  | yes |
| EMX2OS |  | yes |
| ENHO |  | yes |
| ENKUR |  | yes |
| EOMES | yes |  |
| EPHB1 |  | yes |
| EPHX1 |  | yes |
| ERBB2 | yes |  |
| EYA1 |  | yes |
| F2R | yes |  |
| F3 |  | yes |
| FABP5 | yes |  |
| FADS2 |  | yes |
| FAM116B | yes |  |
| FAM158A | yes |  |
| FAM167A |  | yes |
| FAM171B |  | yes |
| FAM181A | yes |  |
| FAM189A2 |  | yes |
| FAM211A |  | yes |
| FAM59B |  | yes |
| FAM64A | yes |  |
| FAM89B | yes |  |
| FAT1 |  | yes |
| FBLN1 |  | yes |
| FBXO2 |  | yes |
| FEZF2 | yes |  |
| FGF2 |  | yes |
| FGFR3 |  | yes |
| FGFRL1 | yes |  |
| FHOD3 | yes |  |
| FJX1 |  | yes |
| FKBP10 | yes |  |
| FLJ42875 | yes |  |
| FLJ46906 | yes |  |
| FOXM1 | yes |  |
| FOXP4 | yes |  |
| FREM2 |  | yes |
| FXYD1 |  | yes |
| FZD2 | yes |  |
| FZD7 | yes |  |
| FZD8 | yes |  |
| GABRA2 |  | yes |
| GABRG1 |  | yes |
| GDPD2 |  | yes |
| GFAP |  | yes |
| GINS2 | yes |  |
| GJA1 |  | yes |
| GJB2 |  | yes |
| GJB6 |  | yes |
| GLI2 | yes |  |
| GLI3 | yes |  |
| GLIS3 |  | yes |
| GNA14 |  | yes |
| GPAM |  | yes |
| GPC1 | yes |  |
| GPC4 | yes |  |
| GPC5 |  | yes |
| GPR125 |  | yes |
| GPR143 |  | yes |
| GPR37L1 |  | yes |
| GPR75-ASB3 |  | yes |
| GPR98 |  | yes |
| GPSM1 | yes |  |
| GPT2 |  | yes |
| GRAMD1C |  | yes |
| GRIK5 | yes |  |
| GRIN2C |  | yes |
| GSG2 | yes |  |
| GSTM5 |  | yes |
| GTSE1 | yes |  |
| H1FX | yes |  |
| HELLS | yes |  |
| HEPACAM |  | yes |
| HEPH |  | yes |
| HES5 | yes |  |
| HES6 | yes |  |
| HHATL |  | yes |
| HIST1H1A | yes |  |
| HIST1H1B | yes |  |
| HIST1H1D | yes |  |
| HIST1H1E | yes |  |
| HIST1H2AC | yes |  |
| HIST1H2AG | yes |  |
| HIST1H2AH | yes |  |
| HIST1H2AI | yes |  |
| HIST1H2AJ | yes |  |
| HIST1H2AK | yes |  |
| HIST1H2AL | yes |  |
| HIST1H2AM | yes |  |
| HIST1H2BB | yes |  |
| HIST1H2BF | yes |  |
| HIST1H2BH | yes |  |
| HIST1H2BJ | yes |  |
| HIST1H2BL | yes |  |
| HIST1H2BM | yes |  |
| HIST1H2BO | yes |  |
| HIST1H3A | yes |  |
| HIST1H3B | yes |  |
| HIST1H3C | yes |  |
| HIST1H3E | yes |  |
| HIST1H3F | yes |  |
| HIST1H3H | yes |  |
| HIST1H3I | yes |  |
| HIST1H3J | yes |  |
| HIST2H2AA3 | yes |  |
| HIST2H2AB | yes |  |
| HIST2H2AC | yes |  |
| HJURP | yes |  |
| HMGN5 | yes |  |
| HPR |  | yes |
| HPSE2 |  | yes |
| HSD17B6 |  | yes |
| HSPB1 | yes |  |
| HSPB8 |  | yes |
| ID4 |  | yes |
| IDI2-AS1 |  | yes |
| IGDCC4 | yes |  |
| IGF2BP2 | yes |  |
| IGF2BP3 | yes |  |
| IGFBP2 | yes |  |
| IL33 |  | yes |
| INHBB | yes |  |
| INSM1 | yes |  |
| INTU | yes |  |
| IQCA1 |  | yes |
| ITGA7 |  | yes |
| KAL1 |  | yes |
| KCNN3 |  | yes |
| KCNQ2 | yes |  |
| KCTD15 | yes |  |
| KIAA0101 | yes |  |
| KIAA0195 |  | yes |
| KIAA1161 |  | yes |
| KIF11 | yes |  |
| KIF15 | yes |  |
| KIF18A | yes |  |
| KIF18B | yes |  |
| KIF20A | yes |  |
| KIF23 | yes |  |
| KIF24 | yes |  |
| KIF2C | yes |  |
| KIF4A | yes |  |
| KREMEN1 |  | yes |
| LAMA1 | yes |  |
| LFNG |  | yes |
| LGI4 |  | yes |
| LGR4 |  | yes |
| LHX2 | yes |  |
| LINC00340 | yes |  |
| LIX1 |  | yes |
| LLGL1 | yes |  |
| LMNA | yes |  |
| LMNB1 | yes |  |
| LMNB2 | yes |  |
| LOC100129148 | yes |  |
| LOC100129518 |  | yes |
| LOC100130522 | yes |  |
| LOC100130894 |  | yes |
| LOC100134868 | yes |  |
| LOC100144602 | yes |  |
| LOC100192426 | yes |  |
| LOC100289473 |  | yes |
| LOC100505659 | yes |  |
| LOC100505875 | yes |  |
| LOC100506795 |  | yes |
| LOC100507206 | yes |  |
| LOC100652999 | yes |  |
| LOC285441 |  | yes |
| LOC439950 | yes |  |
| LOC649395 | yes |  |
| LRIG1 |  | yes |
| LRRC16A |  | yes |
| LRRC37A4 | yes |  |
| LRRC3B |  | yes |
| LTBP1 | yes |  |
| MAP3K12 | yes |  |
| MAPK4 |  | yes |
| MCM10 | yes |  |
| MDK | yes |  |
| MEIS2 | yes |  |
| MELK | yes |  |
| MEX3A | yes |  |
| MFAP2 | yes |  |
| MFAP4 | yes |  |
| MFSD10 | yes |  |
| MGST1 |  | yes |
| MID1 | yes |  |
| MKI67 | yes |  |
| MLC1 |  | yes |
| MMD2 |  | yes |
| MMP28 |  | yes |
| MN1 | yes |  |
| MOK | yes |  |
| MOXD1 | yes |  |
| MRO |  | yes |
| MRVI1 |  | yes |
| MSI1 | yes |  |
| MSMB | yes |  |
| MT1G |  | yes |
| MT3 |  | yes |
| MTRNR2L1 | yes |  |
| MTSS1L | yes |  |
| MTX1 | yes |  |
| MUC1 | yes |  |
| MVD | yes |  |
| MYBL2 | yes |  |
| MYBPC1 |  | yes |
| MZT2A | yes |  |
| NAT8L |  | yes |
| NBPF10 | yes |  |
| NCAN |  | yes |
| NCAPG | yes |  |
| NCAPH | yes |  |
| NDC80 | yes |  |
| NEK2 | yes |  |
| NEUROD2 | yes |  |
| NFATC4 |  | yes |
| NHLH1 | yes |  |
| NIT1 | yes |  |
| NKAIN3 |  | yes |
| NKAIN4 | yes |  |
| NME1-NME2 | yes |  |
| NME3 | yes |  |
| NME7 | yes |  |
| NNAT | yes |  |
| NOTCH1 | yes |  |
| NR2E1 | yes |  |
| NR2F1 | yes |  |
| NRG1 | yes |  |
| NT5DC2 | yes |  |
| NTM |  | yes |
| NTRK2 |  | yes |
| NTSR2 |  | yes |
| NUF2 | yes |  |
| NUSAP1 | yes |  |
| NWD1 |  | yes |
| OAF |  | yes |
| ODF3L1 | yes |  |
| OLFM2 |  | yes |
| OTX1 | yes |  |
| P2RY1 |  | yes |
| PAFAH1B3 | yes |  |
| PALLD | yes |  |
| PALM | yes |  |
| PAMR1 |  | yes |
| PAPLN |  | yes |
| PAX6 | yes |  |
| PBXIP1 |  | yes |
| PCDHGA12 | yes |  |
| PCDHGA3 |  | yes |
| PCDHGC5 | yes |  |
| PDK2 |  | yes |
| PDLIM3 | yes |  |
| PEMT | yes |  |
| PGAP2 | yes |  |
| PHF21B | yes |  |
| PHKA1 |  | yes |
| PHYHD1 |  | yes |
| PIPOX |  | yes |
| PLA2G5 |  | yes |
| PLCD1 |  | yes |
| PLEKHA7 | yes |  |
| PLIN5 |  | yes |
| PLK1 | yes |  |
| PLK4 | yes |  |
| PLSCR4 |  | yes |
| PLTP |  | yes |
| PLXNA3 | yes |  |
| PLXNB1 |  | yes |
| POU3F4 | yes |  |
| PPAP2B |  | yes |
| PPDPF | yes |  |
| PPP1R17 | yes |  |
| PPP1R1B |  | yes |
| PPP1R3C |  | yes |
| PPP1R3G |  | yes |
| PRC1 | yes |  |
| PRDM16 | yes |  |
| PRKG1 |  | yes |
| PRODH |  | yes |
| PRRT2 | yes |  |
| PRSS35 |  | yes |
| PSD2 |  | yes |
| PTCHD1 |  | yes |
| PTMS | yes |  |
| PTPRF | yes |  |
| PTPRS | yes |  |
| PTX3 | yes |  |
| PYCR1 | yes |  |
| PYGM |  | yes |
| RAB36 | yes |  |
| RAD54L | yes |  |
| RAD9A | yes |  |
| RANBP3L |  | yes |
| RBM14-RBM4 | yes |  |
| RFC4 | yes |  |
| RFX2 | yes |  |
| RFX4 |  | yes |
| RGMA | yes |  |
| RGR |  | yes |
| RGS20 |  | yes |
| RLBP1 |  | yes |
| RNF182 |  | yes |
| RNF43 |  | yes |
| RORB |  | yes |
| RPE65 |  | yes |
| RPL13AP5 | yes |  |
| RPL18A | yes |  |
| RPPH1 | yes |  |
| RPS15 | yes |  |
| RPS28 | yes |  |
| RRM2 | yes |  |
| RYR3 |  | yes |
| S100A1 |  | yes |
| SAPCD1 | yes |  |
| SBK1 | yes |  |
| SCARA3 |  | yes |
| SCARNA27 | yes |  |
| SCARNA3 | yes |  |
| SCUBE1 | yes |  |
| SDC2 |  | yes |
| SDC4 |  | yes |
| SDS |  | yes |
| SELENBP1 |  | yes |
| SEMA3A | yes |  |
| SEMA4A |  | yes |
| SEMA4B |  | yes |
| SEMA5B | yes |  |
| SERPINE2 |  | yes |
| SFRP2 | yes |  |
| SFXN5 |  | yes |
| SGK223 | yes |  |
| SGOL1 | yes |  |
| SHROOM3 | yes |  |
| SKA1 | yes |  |
| SKA3 | yes |  |
| SLC13A5 |  | yes |
| SLC14A1 |  | yes |
| SLC16A9 |  | yes |
| SLC1A2 |  | yes |
| SLC1A3 |  | yes |
| SLC1A4 |  | yes |
| SLC25A18 |  | yes |
| SLC25A48 |  | yes |
| SLC39A12 |  | yes |
| SLC44A3 |  | yes |
| SLC4A4 |  | yes |
| SLC6A11 |  | yes |
| SLC7A10 |  | yes |
| SLC7A11 |  | yes |
| SLC7A2 |  | yes |
| SLC9A3R1 |  | yes |
| SLCO1C1 |  | yes |
| SMO | yes |  |
| SNCAIP | yes |  |
| SNTA1 |  | yes |
| SNURF | yes |  |
| SORCS2 |  | yes |
| SOX11 | yes |  |
| SOX2 | yes |  |
| SOX9 | yes |  |
| SPARCL1 |  | yes |
| SPON1 |  | yes |
| ST8SIA2 | yes |  |
| STON2 |  | yes |
| STOX1 |  | yes |
| SYNE2 | yes |  |
| SYPL2 |  | yes |
| TACC3 | yes |  |
| TCF3 | yes |  |
| TCF7L1 | yes |  |
| TFAP2C | yes |  |
| THAP7 | yes |  |
| TKTL1 | yes |  |
| TLCD1 |  | yes |
| TMEM132A | yes |  |
| TMEM158 | yes |  |
| TMEM47 |  | yes |
| TMPRSS3 |  | yes |
| TMSB15A | yes |  |
| TNC | yes |  |
| TNFRSF11B | yes |  |
| TNFSF13 |  | yes |
| TOP2A | yes |  |
| TPD52L1 |  | yes |
| TPX2 | yes |  |
| TRAF7 | yes |  |
| TRIL |  | yes |
| TROAP | yes |  |
| TRPM3 |  | yes |
| TSPAN18 | yes |  |
| TTYH1 |  | yes |
| TUBB2B | yes |  |
| TUBB3 | yes |  |
| UBE2C | yes |  |
| UBE2S | yes |  |
| UG0898H09 | yes |  |
| VAV3 |  | yes |
| VEPH1 | yes |  |
| WEE1 | yes |  |
| WFS1 |  | yes |
| WIF1 |  | yes |
| WNT7B |  | yes |
| WWC1 |  | yes |
| YAP1 |  | yes |
| ZIC2 |  | yes |
| ZIC5 |  | yes |
| ZNF423 | yes |  |
| ZNF663 | yes |  |
| ZNRF3 |  | yes |
