## Supplemental Table 2 for "Computational Identification of Ligand-Receptor Pairs that Drive Human Astrocyte Development"

Panel of 100 targeted RNA-seq genes listed by category (astrocyte-specific, neuronal-specific, reactive, or housekeeping).

| **Gene name** | **Category** |
| --- | --- |
| AGT | Astrocyte |
| ALDH1L1 | Astrocyte |
| AMIGO2 | Reactive |
| APOE | Astrocyte |
| AQP4 | Astrocyte |
| ATP1A2 | Astrocyte |
| BCL11B | Neuron |
| BMPR1B | Astrocyte |
| C3 | Reactive |
| CALB1 | Neuron |
| CD44 | Astrocyte |
| CDH8 | Neuron |
| CDON | Astrocyte |
| CLU | Astrocyte |
| CNTNAP2 | Neuron |
| CRYAB | Astrocyte |
| CXCL1 | Reactive |
| CXCL10 | Reactive |
| DCX | Neuron |
| DIO2 | Astrocyte |
| DLX1 | Neuron |
| DLX2 | Neuron |
| EDNRB | Astrocyte |
| EGFR | Astrocyte |
| EMX1 | Neuron |
| ENPP6 | Astrocyte |
| EOMES | Neuron |
| FABP7 | Neuron |
| FABP7 | Astrocyte |
| FAM107A | Neuron |
| FBRSL1 | Housekeeping |
| FGFR3 | Astrocyte |
| FOXG1 | Neuron |
| GAD1 | Neuron |
| GFAP | Astrocyte |
| GJA1 | Astrocyte |
| GJB7 | Neuron |
| GRIA4 | Neuron |
| GRIN1 | Neuron |
| GRIN2A | Neuron |
| GRM1 | Neuron |
| HOPX | Neuron |
| ID4 | Astrocyte |
| ITGA7 | Astrocyte |
| KCNJ3 | Neuron |
| KIAA0586 | Housekeeping |
| LCN2 | Reactive |
| LHX6 | Neuron |
| LRIG1 | Neuron |
| MAP3K2 | Housekeeping |
| METTL26 | Housekeeping |
| MLC1 | Astrocyte |
| NES | Neuron |
| NEUROD2 | Neuron |
| NEUROD4 | Neuron |
| NEUROD6 | Neuron |
| NFASC | Astrocyte |
| NKX2-1 | Neuron |
| NKX2-2 | Neuron |
| OTX1 | Astrocyte |
| PDLIM3 | Astrocyte |
| PLD5 | Neuron |
| PPIE | Housekeeping |
| PPIL2 | Housekeeping |
| PRODH | Astrocyte |
| PTPRZ1 | Neuron |
| RELN | Neuron |
| RFX1 | Housekeeping |
| RYR3 | Astrocyte |
| S100A10 | Reactive |
| S100B | Astrocyte |
| SERPING1 | Reactive |
| SFRP2 | Astrocyte |
| SLC17A6 | Neuron |
| SLC1A2 | Astrocyte |
| SLC1A3 | Astrocyte |
| SLC32A1 | Neuron |
| SLC4A4 | Astrocyte |
| SLC7A10 | Astrocyte |
| SNAP25 | Neuron |
| SOX9 | Astrocyte |
| SOX9 | Astrocyte |
| SP8 | Neuron |
| SPARC | Astrocyte |
| SPARCL1 | Astrocyte |
| SRGN | Reactive |
| STAT3 | Reactive |
| STMN2 | Neuron |
| SYN1 | Neuron |
| SYT1 | Neuron |
| TBR1 | Neuron |
| THAP3 | Housekeeping |
| TNC | Astrocyte |
| TTYH1 | Astrocyte |
| TUBB3 | Neuron |
| VCAN | Reactive |
| VIM | Astrocyte |
| WIF1 | Astrocyte |
| ZBTB22 | Housekeeping |
| ZNF446 | Housekeeping |
