## Supplemental Table 3 for "Computational Identification of Ligand-Receptor Pairs that Drive Human Astrocyte Development"

Set of 25 astrocyte-enriched and 25 neuronal-enriched genes used as a gene signature to assess transcriptomic changes in these populations. Markers derived from human cell type-specific enrichment data in Zhang and Sloan et al. 2016.

| **Neuronal markers** | **Astrocyte markers** |
| --- | --- |
| BCL11B | CRYAB |
| CALB1 | IL33 |
| CBLN2 | GFAP |
| CDH8 | ITGA7 |
| DCX | TNC |
| EOMES | ALDH1L1 |
| GJB7 | SLC1A3 |
| GRIA4 | VIM |
| GRIK1 | NCAN |
| GRIN2A | CDON |
| GRM1 | ID4 |
| KCNJ3 | SOX9 |
| LHX6 | IGFBPL1 |
| NDNF | PDLIM3 |
| NES | ASPM |
| NEUROD4 | SLC1A2 |
| NEUROG2 | SFRP2 |
| PLD5 | AGT |
| PVALB | OTX1 |
| SP8 | GJA1 |
| STMN2 | FABP7 |
| SYT10 | PAX6 |
| TBR1 | AQP4 |
| TRH | FGFR3 |
| TUBB3 | DIO2 |
