## Supplemental Table 4 for "Computational Identification of Ligand-Receptor Pairs that Drive Human Astrocyte Development"

Ligand concentrations used for exposure experiments. Ligands were added every other day to maintain levels throughout culture windows.

| **Ligands** | **ng/ul** |  |
| --- | --- | --- |
| APP | 100 | Ligands 1-5 |
| APOE | 500 |  |
| GAS6 | 250 |  |
| CALR | 200 |  |
| IGF1 | 100 |  |
| TGFB2 | 20 | Ligands 6-10 |
| NLGN1 | 300 |  |
| TSLP | 100 |  |
| DKK1 | 100 |  |
| BMP4 | 20 |  |
